# Evolutionary pathways to antibiotic resistance are dependent upon environmental structure and bacterial lifestyle

**DOI:** 10.1101/581611

**Authors:** Alfonso Santos-Lopez, Christopher W. Marshall, Michelle R. Scribner, Daniel Snyder, Vaughn S. Cooper

**Author notes:** A.S-L. and C.W.M. contributed equally to this manuscript.

## Abstract

Bacterial populations vary in their stress tolerance and population structure depending upon whether growth occurs in well-mixed or structured environments. We hypothesized that evolution in biofilms would generate greater genetic diversity than well-mixed environments and lead to different pathways of antibiotic resistance. We used experimental evolution and whole genome sequencing to test how the biofilm lifestyle influenced the rate, genetic mechanisms, and pleiotropic effects of resistance to ciprofloxacin in *Acinetobacter baumannii* populations. Both evolutionary dynamics and the identities of mutations differed between lifestyle. Planktonic populations experienced selective sweeps of mutations including the primary topoisomerase drug targets, whereas biofilm-adapted populations acquired mutations in regulators of efflux pumps. An overall trade-off between fitness and resistance level emerged, wherein biofilm-adapted clones were less resistant than planktonic but more fit in the absence of drug. However, biofilm populations developed collateral sensitivity to cephalosporins, demonstrating the clinical relevance of lifestyle on the evolution of resistance.

## Introduction

Antimicrobial resistance (AMR) is one of the main challenges facing modern medicine. The emergence and rapid dissemination of resistant bacteria is decreasing the effectiveness of antibiotics and it is estimated that 700,000 people die per year due to AMR-related problems (O’Neill, 2016). AMR, like all phenotypes, is an evolved property, either the ancient product of living amidst other microbial producers of antimicrobials (Martínez, 2008), or the recent product of strong selection by human activities for novel resistance-generating mutations (Ventola, 2015).

The dominant mode of growth for most microbes is on surfaces, and this biofilm lifestyle is central to AMR (Ahmed, Porse, Sommer, Hoiby, & Ciofu, 2018; Hoiby, Bjarnsholt, Givskov, Molin, & Ciofu, 2010; Olsen, 2015), especially in chronic infections (Wolcott, 2017; Wolcott et al., 2010). However, with few exceptions (Ahmed et al., 2018; France, Cornea, Kehlet-Delgado, & Forney, 2019; Ridenhour et al., 2017), most of the research on the evolution of AMR has been conducted in well-mixed populations [reviewed in (Hughes & Andersson, 2017)] or on agar plates (Michael Baym et al., 2016), conditions that cannot simulate the effects of biofilms on the evolution of AMR. Consequently, our understanding of how this lifestyle influences the evolution of AMR, whether by different population-genetic dynamics or molecular mechanisms, is limited. One example is that the close proximity of cells in biofilms may facilitate the horizontal transfer and persistence of resistance genes in bacterial populations (Ridenhour et al., 2017; Stalder & Top, 2016). Less appreciated is the potential for the biofilm lifestyle to influence the evolution of AMR by *de novo* chromosomal mutations. This emergence of AMR in biofilms is important because: i) the environmental structure of biofilms can increase clonal interference, rendering selection less effective and enhancing genetic diversity (Cooper, Staples, Traverse, & Ellis, 2014; Ellis, Traverse, Mayo-Smith, Buskirk, & Cooper, 2015; France et al., 2019; Habets, Rozen, Hoekstra, & de Visser, 2006; Traverse, Mayo-Smith, Poltak, & Cooper, 2013), ii) distinct ecological conditions within the biofilm can favor functionally distinct adaptations to different niches (Poltak & Cooper, 2011), iii) the biofilm itself can protect its residents from being exposed to external stresses like antibiotics or host immunity (Eze, Chenia, & El Zowalaty, 2018; E. Geisinger & Isberg, 2015), and iv) slower growth within biofilms can reduce the efficacy of antibiotics that preferentially attack fast-growing cells (Kirby, Garner, & Levin, 2012; Walters, Roe, Bugnicourt, Franklin, & Stewart, 2003). The first two hypotheses would predict more complex evolutionary dynamics within biofilms than in well-mixed environments (Steenackers, Parijs, Foster, & Vanderleyden, 2016), while the second two predict different rates of evolution, targets of selection, and likely less potent mechanisms of AMR (Andersson & Hughes, 2014). Together, these potential factors call into question the conventional wisdom of a tradeoff between fitness and antimicrobial resistance, a relationship that remains to be clearly defined.

Here, we study the evolutionary dynamics and effects of new resistance mutations in the opportunistic nosocomial pathogen *Acinetobacter baumannii*, which is often intrinsically resistant to antibiotics or has been reported to rapidly evolve resistance to them (Doi, Murray, & Peleg, 2015). This pathogen is categorized as one of the highest threats to patient safety (Asif, Alvi, & Rehman, 2018), partly due to its ability to live on inanimate surfaces in biofilms (Eze et al., 2018). We experimentally propagated populations of *A. baumannii* exposed either to subinhibitory or increasing concentrations of ciprofloxacin (CIP) over 80 generations in biofilm or planktonic conditions to ascertain whether these lifestyles select for different mechanisms of AMR. Rather than focusing on the genotypes of single isolates, which can limit the scope of an analysis, we conducted whole-population genomic sequencing over time to define the dynamics of adaptation and the fitness of certain resistance alleles compared to others in the experiment. We then identified clones with specific genotypes that we linked to fitness and resistance phenotypes. This approach sheds new light on the ways that pathogens adapt to antibiotics while growing in biofilms and has implications for treatment decisions.

## Results and Discussion

### 1. Experimental evolution

Replicate cultures of the susceptible *A. baumannii* strain ATCC 17978 (Baumann, Doudoroff, & Stanier, 1968; Piechaud & Second, 1951) were established under planktonic or biofilm conditions in one of three treatments: i) no antibiotics, ii) sub-inhibitory concentration of the antibiotic ciprofloxacin (CIP) and iii) evolutionary rescue (Bell & Gonzalez, 2009) in which CIP concentrations were increased every 72 hours from subinhibitory concentrations to four times the minimum inhibitory concentration (MIC) (Figure 1A). Before the start of the antibiotic evolution experiment, we propagated the ATCC strain for ten days in planktonic conditions to reduce the influence of adaptation to the laboratory conditions on subsequent comparisons. CIP was chosen because of its clinical importance in treating *A. baumannii* (Ardebili, Lari, & Talebi, 2014; Doi et al., 2015; Lopes & Amyes, 2013), its ability to penetrate the biofilm matrix (Tseng et al., 2013) allowing similar efficacy in well mixed and structured populations (Kirby et al., 2012), and because it is not known to stimulate biofilm formation in *A. baumannii* (Aka & Haji, 2015). Planktonic populations were serially passaged by daily 1:100 dilution while biofilm populations were propagated using a bead model simulating the biofilm life cycle (Poltak & Cooper, 2011; Traverse et al., 2013; Turner, Marshall, & Cooper, 2018). This model selects for bacteria that attach to a 7 mm polystyrene bead, form a biofilm, and then disperse to colonize a new bead each day. (A video tutorial for this protocol is available at http://evolvingstem.org). The transfer population size in biofilm and in planktonic cultures was set to be nearly equivalent at the beginning of the experiment (approximately 1×10^7^ CFU/mL), because population size influences mutation availability and the response to selection (Cooper, 2018; Salverda, Koomen, Koopmanschap, Zwart, & de Visser, 2017). The mutational dynamics of three lineages from each treatment were tracked by whole-population genomic sequencing (Figure 1A). We also sequenced 49 single clones isolated from 22 populations at the end of the 12-day experiment to determine mutation linkage.

**Figure 1.**
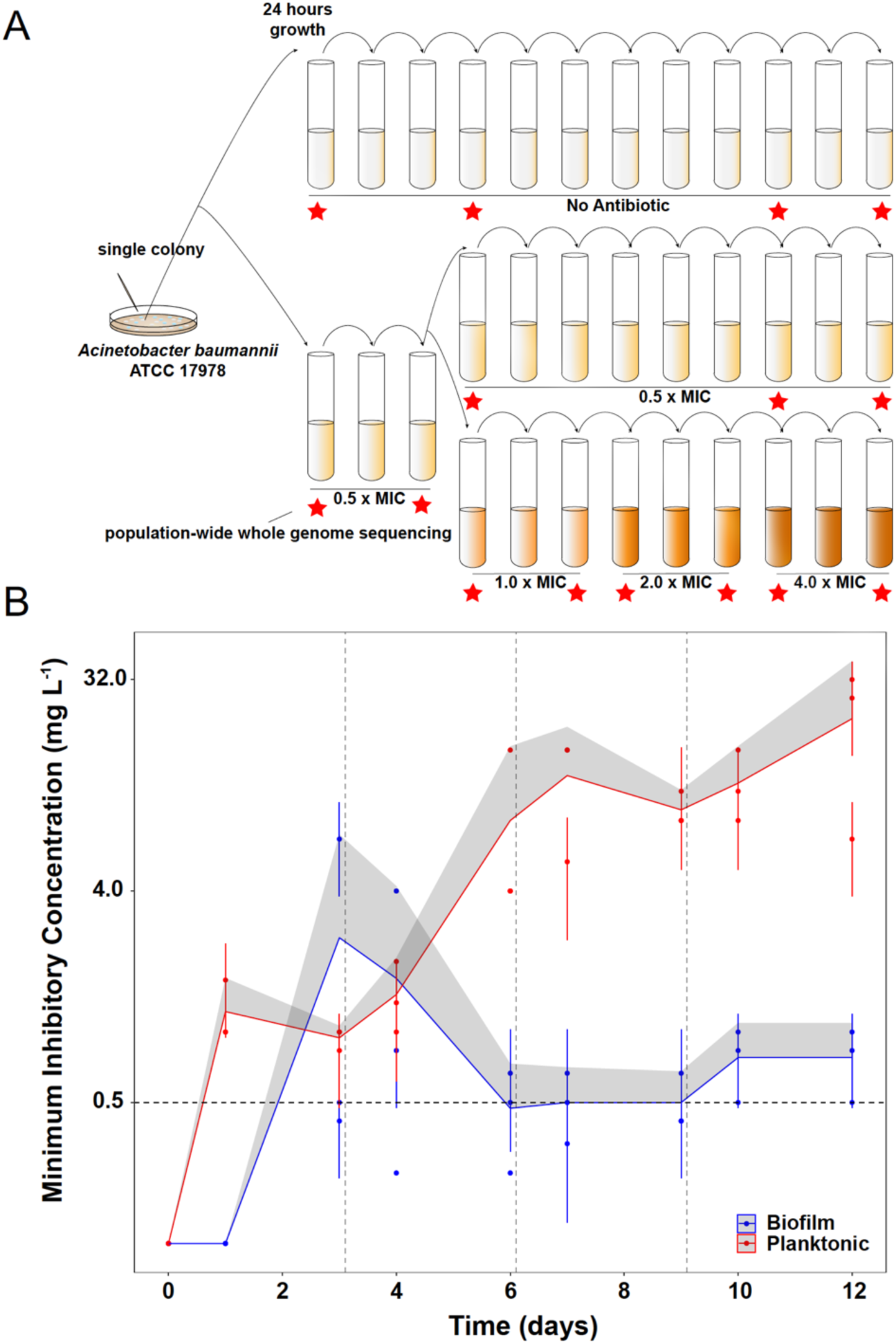
Experimental design (A) and dynamics of evolved resistance levels during the evolutionary rescue experiment (B). A) A single clone of *A. baumannii* ATCC 17978 was propagated both in biofilm and planktonic conditions for 12 days under no antibiotics (top), subinhibitory concentrations of CIP (0.0625 mg/L = 0.5x MIC) (middle) or in increasing concentrations of CIP (bottom). For the latter, termed evolutionary rescue, the concentration of CIP was doubled from 0.5 × MIC to 4.0 × MIC every 72 hours. As a control, five populations of *A. baumannii* ATCC 17978 were propagated in biofilm and five in planktonic conditions in the absence of antibiotics. We estimated the MICs to CIP and froze the populations for sequencing before and after doubling the antibiotic concentrations (red stars). B) MICs (mg/L) of CIP were measured for replicate populations during the evolutionary rescue. The red and blue points represent the MICs of three populations propagated in planktonic or biofilm, respectively, with the 95% CI represented by the error bars. The red and blue lines represent the grand mean of the three planktonic and biofilm populations, respectively, with the upper 95% CI indicated by the grey shaded area. Horizontal dashed line indicates the highest CIP exposure during the experiment (4x MIC) and vertical lines indicate time when CIP concentration was doubled.

### 2. Evolution of CIP resistance

The rate and extent of evolved resistance depends on the strength of antibiotic selection (Andersson & Hughes, 2014; Oz et al., 2014), the distribution of fitness effects of mutations that increase resistance to the drug (Maclean, Hall, Perron, & Buckling, 2010), and the population size of replicating bacteria (Cooper, 2018; Salverda et al., 2017). The mode of bacterial growth can in principle alter each of these three variables and generate different dynamics and magnitudes of AMR. In the populations exposed to the increasing concentrations of CIP (the evolutionary rescue), the magnitude of evolved CIP resistance differed between planktonic and biofilm populations. Planktonic populations became approximately 160x more resistant on average than the ancestral clone while the biofilm populations became only 6x more resistant (Figure 1B and Table S1). Planktonic populations also evolved resistance much more rapidly, becoming 10x more resistant after only 24 hours of growth in sub-inhibitory CIP. This level of resistance would have been sufficient for surviving the remainder of the experiment, but MICs continued to increase at each sampling (Figure 1B). The evolution of resistance far beyond the selective requirement indicates that mutations conferring higher resistance also increased fitness in planktonic populations exposed to CIP.

In contrast, biofilm-evolved populations evolved under the evolutionary rescue regime acquired much lower levels of resistance (*ca.* 3– 7x the ancestral MIC) and primarily in a single step between days 3 and 4 (Figure 1B). In one notable exception, the MIC of biofilm population B2 increased *∼*50x after 3 days of selection in subinhibitory concentrations of CIP (Figure 1B), but then the resistance of this population declined to only 6x higher than the ancestral strain. This dynamic suggested that a mutant conferring high-level resistance rose to intermediate frequency but was replaced by a more fit, yet less resistant, mutant (this possibility is evaluated below).

Lower levels of resistance were observed in populations selected at constant subinhibitory concentrations of CIP. Biofilm populations were 4x more resistant than the ancestor and planktonic populations were 20x more resistant (Table S1). We can infer that biofilm growth does not select for the high-level resistance seen in planktonic populations, instead favoring mutants with low levels of resistance and better adapted to life in a biofilm. It is important to note that these MIC measurements were made in planktonic conditions according to the clinical standards (CLSI, 2007) and that these values increased when measured in biofilm (Table S2). Our results correspond with studies of clinical isolates in which those producing more biofilm (and likely having adapted in biofilm conditions) were less resistant than non-biofilm-forming isolates (Wang et al., 2018). Nevertheless, measuring growth and MIC is context-dependent (Borriello et al., 2004; Hill et al., 2005; Kirby et al., 2012), and because the biofilm environment at least partially protects cells from antibiotic exposure (Table S2), it can be argued that evolved biofilm populations experienced lower CIP concentrations than planktonic populations. However, we selected CIP because it can penetrate the biofilm barrier (Tseng et al., 2013), and furthermore, cells growing in the bead model must disperse from one bead to colonize the next one in a less protected state. Overall, these results demonstrate that exposing bacteria to low levels of antibiotic risks selection for high levels of resistance that can make future treatment more difficult (Wistrand-Yuen et al., 2018).

### 3. Evolutionary dynamics under CIP treatment

In large bacterial populations (>10^5^ cells) growing under strong selection, adaptive mutations conferring beneficial traits such as antibiotic resistance will dominate population dynamics (Jeffrey E Barrick & Lenski, 2013; Cooper, 2018). Therefore, if a single mutation renders the antibiotic ineffective and provides the highest fitness gain, it would be expected to outcompete all other less fit mutations. Further, the stronger the selection for resistance, the greater the probability of genetic parallelism among replicate populations (Bolnick, Barrett, Oke, Rennison, & Stuart, 2018). Under the conditions of these experiments, approximately 10^6^ mutations occur in the first growth cycle and roughly 10^7^ mutations arise over the 12 days of selection, leading to a probability of 0.98 that every site in the 4Mbp *A. baumannii* genome experiences a mutation at least once over the course of the 12-day experiment (see Table S3 for details of these calculations). The dramatic differences in the evolved resistance levels of planktonic and biofilm populations suggested distinct genetic causes of resistance resulting from different selective forces in these treatments. We also predicted greater genetic diversity in the biofilm treatments, owing to spatial structure and/or niche differentiation (Traverse et al., 2013), than in the planktonic cultures, in which we expected selective sweeps (Jeffrey E. Barrick et al., 2009). A signature of spatial structure alone might be different mutations in the same gene with predicted similar function coexisting over time, which is a form of clonal interference (de Visser & Rozen, 2006). A signature of niche differentiation might be the coexistence of mutations in different genes with unique functions, which is a form of adaptive radiation (Kassen, 2009).

We conducted whole-population genomic sequencing of three replicates per treatment to identify all contending mutations above a detection threshold of 5% (see Methods). The spectrum of mutations from CIP-treated populations are consistent with expectations from strong positive selection on altered or disrupted coding sequences (see Table 1 for day-12 results and Table S4 for dynamics across the experiment). High nonsynonymous to synonymous mutation ratios were observed in both lifestyles (8.5 in planktonic and 9.7 in biofilm). 43% of the total mutations in planktonic and 34% in biofilm were insertions or deletions, which is vastly enriched over typical mutation rates of ∼10 SNPs/indel under neutral conditions (Dillon, Sung, Sebra, Lynch, & Cooper, 2017; Lynch et al., 2016). Roughly 30% of the mutations in CIP-treated populations of either lifestyle occurred in intergenic regions (30% in planktonic-propagated populations and 32% in biofilm). Of the intergenic mutations, 72% of the planktonic mutations and 18% of the biofilm mutations occurred in promoters, 5’ untranslated regions, or in putative terminators (Kröger et al., 2018).

**Table 1.**
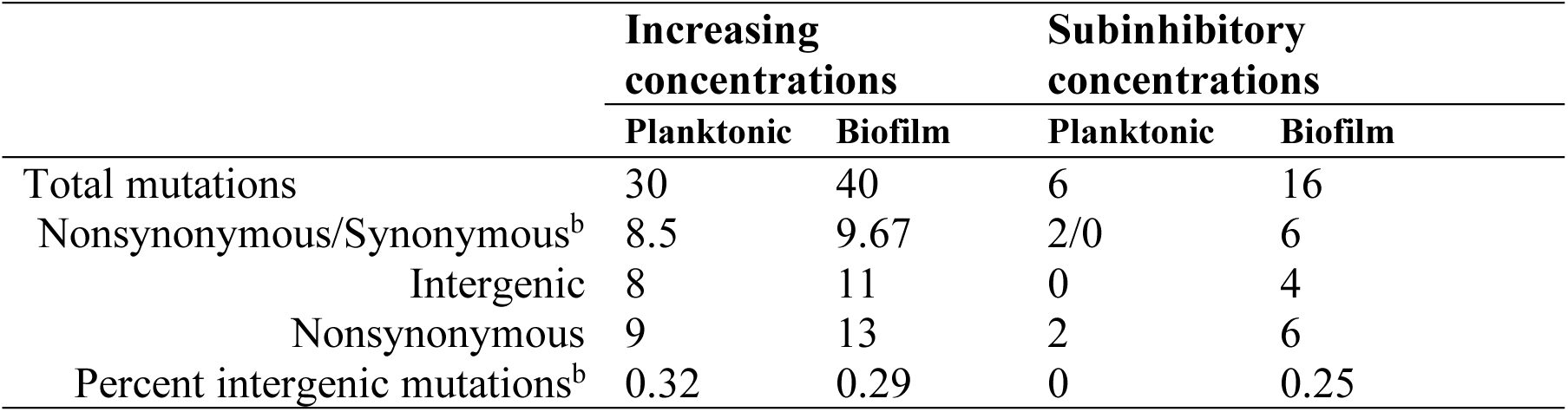
Mutation spectrum of different selective environments. Attributes of the contending mutations during the 12 days of the evolution experiment. ^a^Results from the last day of the experimental evolution. ^b^Accounting for all unique mutations detected after filtering (see methods). For mutation dynamics over time, see Table S3.

As expected from theory, in CIP-selected planktonic populations where selection is most efficient, one or two mutations rapidly outcompeted others and fixed (Figure 2). Selection in biofilms, however, produced fewer selective sweeps and maintained more contending mutations, especially at lower antibiotic concentrations. In one population, multiple mutations in the same locus (*adeL*) rose to high frequency and persisted, which is consistent with the effect of population structure alone. In the other two populations, mutations in different efflux pumps (*adeL, adeS, adeN*) contended during the experiment, which could be explained by population structure or ecological diversification, if these mutations produced different phenotypes. Overall, across all treatments and timepoints, biofilm-adapted populations were significantly more diverse than the planktonic-adapted populations (Shannon index; Kruskal Wallis, chi-squared = 7.723, p = 0.005), particularly at subinhibitory concentrations of CIP (Figure S1A). Notably, increasing drug concentrations eliminated the differences in diversity between treatments (Figure S1B), but the greater diversity in biofilms treated with lower doses generated more diversity for selection to act upon in a changing environment. This higher standing diversity is important when considering dosing and when antibiotic exposure may be low (*e.g.* in the external environment or when bound to tissues) (Baquero, Negri, Morosini, & Blazquez, 1998; Khan, Berglund, Khan, Lindgren, & Fick, 2013) because biofilms with more allelic diversity have a greater chance of survival to drug and immune attack (Fux, Costerton, Stewart, & Stoodley, 2005).

**Figure 2.**
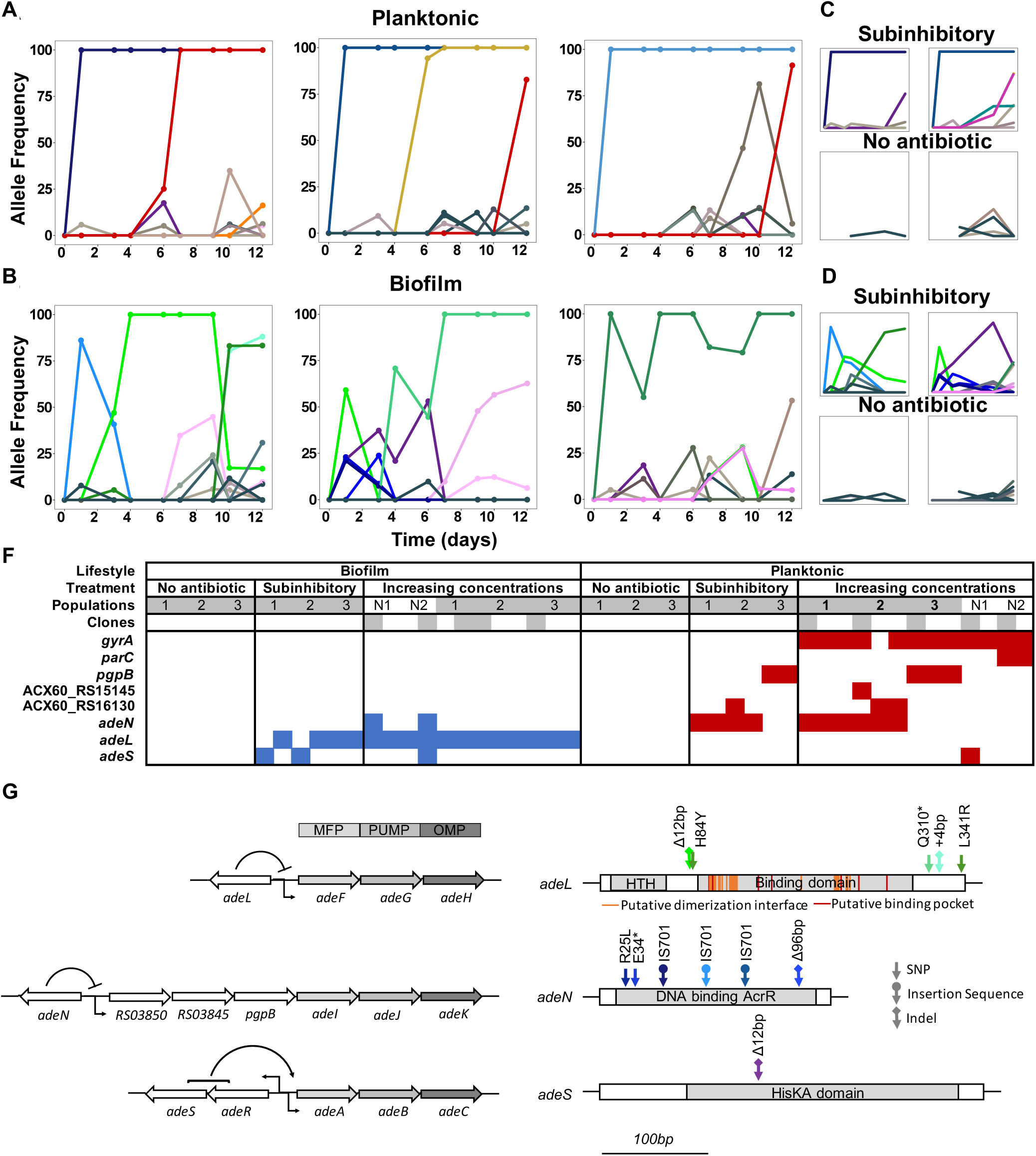
Lifestyle-dependent mutations and dynamics under increasing CIP selection. Mutation frequencies in planktonic (A and C) and biofilm populations (B and D) over time as determined by population-wide whole genome sequencing. A) and B) show the mutation frequencies obtained under increasing concentrations of CIP. From left to right: P1, P2 and P3 in A) and B1, B2 and B3 in B). C) and D) show the mutation frequencies obtained under the subinhibitory (top) and no antibiotic (bottom) treatments. Mutations in the same gene share a color. Blue: *adeN* or genes regulated by *adeN*; green: *adeL*; gold: MFS putative transporter ACX60_RS15145; purple: *adeS*; pink: *sohB*; red: *gyrA;* and orange: *parC*. Grey and brown colors indicate genes potentially unrelated to adaptation to CIP. F) Mutated genes in the sequenced clones. Each column represents one clone. Grey shading of populations indicates whole population sequencing and N1 and N2 indicate populations where only clones were sequenced. Grey shaded clones were used for MIC and fitness estimations. Blue and red indicate SNPs in biofilm and planktonic growing populations respectively. For all SNPs identified in the 49 clones, see Figure S2 and Table S6. G) The genetic organization of the RND efflux pumps is shown on the left. MFP and OMP denote membrane fusion protein and outer membrane protein respectively. All mutations found in the RND regulators are shown on the right.

In contrast with the data observed in the populations evolving under CIP pressure, drug-free control populations contained no mutations that achieved high frequency during the experiment (Figures 2C and 2D). These results suggest that the ancestral starting clone was already well-adapted to our experimental conditions, likely because we had previously propagated the *A. baumannii* ATCC 17978 clone under identical drug-free conditions for 10 days. This preadaptation phase led to the fixation of mutations in three genes (Table S5).

### 4. Lifestyle determines the selected mechanisms of resistance

*A. baumannii* clinical samples acquire resistance to CIP by two principal mechanisms: modification of the direct antibiotic targets — gyrase A or B and topoisomerase IV — or by the overexpression of efflux pumps reducing the intracellular concentrations of the antibiotic (Doi et al., 2015). To directly associate genotypes with resistance phenotypes, we sequenced 49 clones isolated at the end of the experiment, the majority of which were selected to delineate genotypes in the evolutionary rescue populations (Figures 2F and S2).

Both the genetic targets and mutational dynamics of selection in planktonic and biofilm environments differed. Mutations disrupting three negative regulators of efflux pumps evolved in parallel across populations exposed to CIP, but mutations in two of these (*adeL* and *adeS*) were nearly exclusive to biofilm clones (Figure 2F). The most common and highest frequency mutations observed in the biofilm populations were in the repressor gene *adeL* (Figures 2F, S2, and Table S6), which regulates AdeFGH, one of three resistance-nodulation-division (RND) efflux pump systems in *A. baumannii* (Coyne, Rosenfeld, Lambert, Courvalin, & Perichon, 2010; Fernando, Zhanel, & Kumar, 2013; Pournaras, Koumaki, Gennimata, Kouskouni, & Tsakris, 2016). In the planktonic lines, the predominant mutations were found in *adeN,* which is a negative regulator of AdeIJK and were mainly insertions of IS701 that disrupted the gene (X.-Z. Li, C. A. Elkins, & H. I. Zgurskaya, 2016).

In biofilm lines, different contending *adeL* mutations were detected in each replicate after 24 hours then eventually fixed as CIP concentrations increased (green lines in Figure 2B), sometimes along with a secondary *adeL* mutation. This pattern suggests that altering efflux via *adeL* generates adaptations to the combination of CIP and biofilm which is supported by the increase in biofilm formation by the *adeL* mutants (Figure S3). Further, mutants with higher resistance than necessary appear to be maladaptive in the biofilm treatment. For example, *adeN* (found more often in planktonic culture) and *adeS* mutations found simultaneously on day 3 in population B2 (Figure 2) led to a spike in resistance at that timepoint (Figure 1), but these alleles were subsequently outcompeted by *adeL* mutants that were evidently more fit despite lower planktonic resistance.

In contrast to the biofilm populations, all planktonic populations with increasing concentrations of CIP eventually acquired a single high frequency mutation in *gyrA* (S81L), the canonical ciprofloxacin-resistant mutation in DNA gyrase. These *gyrA* mutations evolved in genetic backgrounds containing either an *adeN* mutant or a *pgpB* mutant. *pgpB* is a gene that encodes a putative membrane associated lipid phosphatase and is co-regulated by *adeN* (Hua, Chen, Li, & Yu, 2014). Other mutations associated with high levels of resistance affected *parC,* encoding topoisomerase IV, and regulatory regions of two putative transporters, ACX60_RS15145 and ACX60_RS1613, the latter being co-transcribed with the multidrug efflux pump *abeM* (Su, Chen, Mizushima, Kuroda, & Tsuchiya, 2005). Few other mutations exceeded the 10% of the total population filter in the planktonic lines. The rapid fixation of only *adeN* and *adeN*-regulated alleles in the planktonic CIP-exposed populations indicate that *adeN* conferred higher fitness than other CIP-resistant mutations at low drug concentrations. Subsequently, at increased concentrations of CIP, on-target mutations in *gyrA* were favored in each line.

Together, our results demonstrate that bacterial lifestyle influences the evolutionary dynamics and targets of selection of AMR. Loss-of-function mutations in regulators of the *adeFGH* and *adeABC* RND efflux pumps that increased CIP resistance ∼4-fold in biofilm populations treated with CIP. Adaptation by planktonic populations exposed to CIP proceeded first by altering the *adeN*-controlled *adeIJK* efflux pump and then by directly altering the targets of the fluoroquinolone, *gyrA* and *parC*, leading to much higher levels of resistance.

### 5. Evolutionary consequences of acquiring resistance

The large population sizes (10^7^ – 10^9^ cells) and number of generations (∼80) in all evolved lines mean that similar mutations very likely arose in each replicate regardless of treatment, meaning that the success of some mutations over others reflects their greater fitness in that condition (Table S3) (Cooper, 2018). Yet *de novo* acquired antibiotic resistance is often associated with a fitness cost in the absence of antibiotics [reviewed in (Vogwill & MacLean, 2015)]. The extent of this cost and the ability to compensate for it by secondary mutations (compensatory evolution) is a key attribute determining the spread and maintenance of the resistance mechanism (Maclean et al., 2010; Moore, Rozen, & Lenski, 2000; Vogwill & MacLean, 2015; Zhao & Drlica, 2002). A negative correlation between CIP resistance and fitness of resistant genotypes in the absence of antibiotics has been previously described in *Escherichia coli,* suggesting a trade-off between these traits (Basra et al., 2018; Huseby et al., 2017; Marcusson, Frimodt-Moller, & Hughes, 2009).

To determine the relationship between resistance and fitness in the absence of antibiotics in our experiment, we chose 10 clones (5 each from biofilm and planktonic populations, Figures 2F and S2) with different genotypes and putative resistance mechanisms and measured their resistance and fitness phenotypes in both planktonic and biofilm conditions (Figure 3). As expected from the populations (Figure 1B), the biofilm clones much were less resistant in planktonic conditions than those evolved planktonically [MIC = 0.58 mg/L (SEM = 0.13) vs. MIC = 8.53 mg/L (SEM = 1.96), two-tailed t-test: p < 0.05, t = 4.048, df = 80]. However, biofilm-evolved clones were more fit relative to the ancestral strain than the planktonic-evolved clones in the absence of antibiotic (two-tailed t-test: p = 0.008, t = 2.984 df = 18) (Figure 3). Importantly, these fitness measurements were made in both planktonic and biofilm conditions, demonstrating that even in the conditions they evolved in, the planktonic selected clones were less fit as a result of antibiotic resistance trade-offs. However, one planktonic-evolved clone with mutations in both *gyrA* and *parC* exhibited no significant fitness cost and high levels of resistance. This suggests that, as in *Pseudomonas aeruginosa*, the *parC* mutation may compensate for the cost imposed by *gyrA* mutation (Kugelberg, Lofmark, Wretlind, & Andersson, 2005), an example of sign epistasis (Sackman & Rokyta, 2018). Overall, mutants selected in biofilm-evolved populations were less resistant than mutants selected in planktonic populations (Figure 1B) but produced more biofilm (Figure S3) and paid little or no fitness cost in the absence of antibiotics (Figures 3). This cost-free resistance implies that these subpopulations could persist in the absence of drug, limiting the treatment options and demanding new approaches to treat high fitness, resistant pathogens (Baym, Stone, & Kishony, 2016).

**Figure 3.**
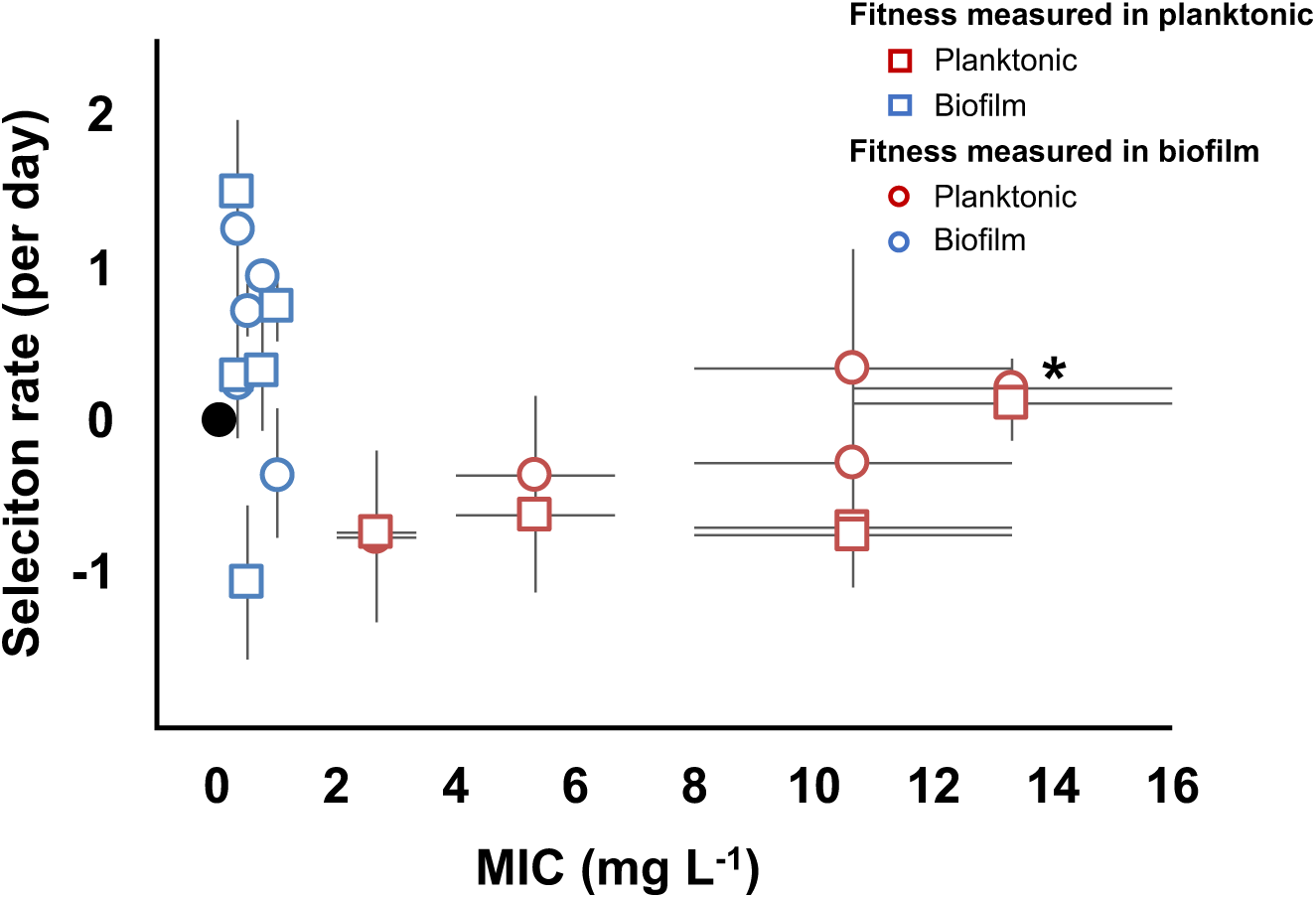
Evolved trade-off between resistance level and fitness. Relative fitness (average ± SEM) of 10 evolved clones from the evolutionary rescue experiment compared to the ancestor and their MICs (mg/L) to CIP. Fitness was measured in both planktonic (squares) and biofilm (circles) conditions. MICs were estimated in planktonic conditions. Black dot represents the ancestral clone. *Denotes the clone with *gyrA* and *parC* mutations.

### 6. Evolutionary interactions with other antibiotics

When a bacterium acquires resistance to one antibiotic, the mechanism of resistance can also confer resistance to other antibiotics (cross-resistance) or increase the susceptibility to other antibiotics (collateral sensitivity) (Pal, Papp, & Lazar, 2015). We tested the MIC of the evolved populations to 23 different antibiotics in planktonic conditions. We observed changes in susceptibilities to 13 of the 23 antibiotics tested, and these changes were growth mode dependent (Figure 4). For example, planktonic populations exhibited cross resistance to cefpodoxime and ceftazidime, but biofilm populations evolved collateral sensitivity to these cephalosporins. Cross-resistance was associated genetically with *adeN, adeS, gyrA* or *pgpB* mutations, and collateral sensitivity was associated with *adeL* mutations. Selection in these environments evidently favors the activation of different efflux pumps or modified targets that have different pleiotropic consequences for multidrug resistance (Podnecky et al., 2018).

**Figure 4.**
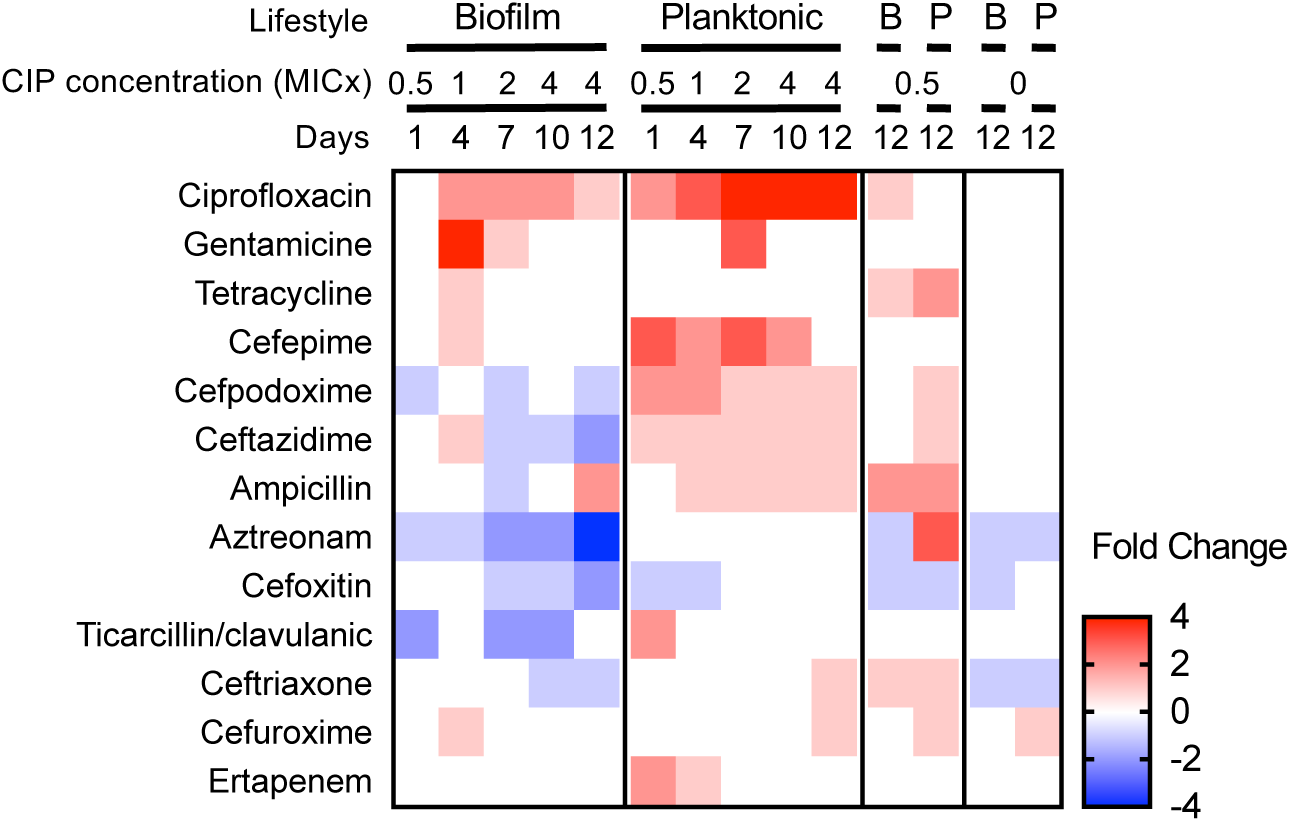
Collateral sensitivities and cross resistances to various antibiotics. Heat map showing the relative changes in antimicrobial susceptibility to 13 of the 23 antibiotics tested in the evolved populations (those not shown had no changes). Results shown are the median values of the fold change in the evolved populations compared to the ancestral strain. For subinhibitory and no-antibiotic treatments, only day 12 is shown.

The mechanisms leading to collateral sensitivity are still poorly understood but they depend on the genetic background of the strain, the nature of the resistance mechanisms (Barbosa et al., 2017; Yen & Papin, 2017), and the specific physiological context of the cells (Leus et al., 2018). In *A. baumannii*, each RND efflux pump is suggested to be specific for certain classes of antibiotics (Coyne, Courvalin, & Perichon, 2011; Leus et al., 2018; X. Z. Li, C. A. Elkins, & H. I. Zgurskaya, 2016). Similar to our results (Figure 4), Yoon and collaborators demonstrated that efflux pumps AdeABC and AdeIJK, regulated by *adeS* and *adeN* respectively, increased the resistance level to some β-lactams when overexpressed (Yoon et al., 2015). However, production of AdeFGH, the efflux pump regulated by *adeL,* decreased resistance to some β-lactams and other families of antibiotics or detergents by an unknown mechanism (Leus et al., 2018; Yoon et al., 2015). Increased sensitivity to β-lactams with efflux overexpression has also been reported in *P. aeruginosa* (Azimi & Rastegar Lari, 2017), which demonstrates the urgency of understanding the physiological basis of collateral sensitivity to control AMR evolution. Exploiting collateral sensitivity has been proposed to counteract the evolution of resistant populations both in bacteria (Imamovic & Sommer, 2013; Kim, Lieberman, & Kishony, 2014; Nichol et al., 2019) and in cancer (Dhawan et al., 2017). Remarkably, our results show that biofilm growth, commonly thought to broaden resistance, may actually generate collateral sensitivity during treatment with CIP and potentially other fluoroquinolones.

### 7. Clinical relevance

Our results demonstrate that the mode of growth determines both the mechanism of evolved resistance and the spectrum of sensitivity to other families of antibiotics. Further, the mutations selected in our experimental conditions also play an important role in clinical isolates, as fluoroquinolone resistance mediated by plasmids in *A. baumannii* appears to be rare (Yang, Hu, Liu, Ye, & Li, 2016). The mutations S81L in *gyrA* and S80L *in parC* acquired by the sensitive ATCC 17978 strain used in this study have been reported worldwide as the primary mechanism conferring high levels of resistance to fluorquinolones in clinical isolates (Adams-Haduch et al., 2008; Dahdouh et al., 2017; W. A. Warner et al., 2016).

In addition to the on-target mechanisms of resistance through gyrase or topoisomerase mutations, *A. baumannii* isolates acquire comparatively moderate levels of fluoroquinolone resistance by modifications in various RND efflux pumps. These RND efflux pumps have overlapping yet differing substrate profiles and may act synergistically in increasing the resistance level (Table S7) (Coyne et al., 2010; Damier-Piolle, Magnet, Bremont, Lambert, & Courvalin, 2008; Fernando et al., 2013; Rosenfeld, Bouchier, Courvalin, & Perichon, 2012). In our experiment, all biofilm and planktonic populations and nearly all isolated clones acquired mutations in at least one of the three regulators of the RND efflux pumps (*adeL, adeS, adeN*) or in a gene regulated by one of these regulators (*pgpB*). Mutations in *adeL* upregulate the expression of the RND efflux pump AdeFGH (Figure 2, Table S7) that is associated with a multidrug resistant phenotype in clinical isolates (Coyne et al., 2010; Fernando et al., 2013; Leus et al., 2018; Pournaras et al., 2016). AdeL-AdeFGH genes are often highly expressed in infection isolates, which could reflect adaptation to the biofilm lifestyle (Coyne et al., 2010; Fernando et al., 2013). Further, overexpression of *adeG* is predicted to enhance transport of acylated homoserine lactones, including quorum-sensing autoinducers, which can increase both drug resistance and biofilm formation (Alav, Sutton, & Rahman, 2018; He et al., 2015). However, in clinical isolates, overexpression of the AdeFGH pump is less common than the AdeIJK efflux pump that is regulated by *adeN* (Rosenfeld et al., 2012; Yoon et al., 2015). AdeIJK contributes to resistance to biocides, hospital disinfectants, and to both intrinsic and acquired antibiotic resistance in *A. baumannii* (Damier-Piolle et al., 2008; Rosenfeld et al., 2012) and may decrease biofilm formation, which could explain its prevalence in planktonic populations here (Yoon et al., 2015). Perhaps more concerning, this study demonstrates that the overexpression of RND efflux pumps as a resistance mechanism may produce little fitness cost in *A. baumannii*, as has previously been demonstrated in both *P. aeruginosa* and *Neisseria gonorrhoeae* (Olivares Pacheco, Alvarez-Ortega, Alcalde Rico, & Martínez, 2017; D. M. Warner, Folster, Shafer, & Jerse, 2007).

## Conclusions

We used experimental evolution of the opportunistic pathogen *A. baumannii* in both well-mixed and biofilm conditions to examine how lifestyle influences the dynamics, diversity, identity of genetic mechanisms, and direct and pleiotropic effects of resistance to a common antibiotic. Experimental evolution is a powerful method of screening naturally arising genetic variation for mutants that are the best fit in a defined condition (Cooper, 2018; Elena & Lenski, 2003; Van den Bergh, Swings, Fauvart, & Michiels, 2018). When population sizes are large and reproductive rates are rapid, as they were here, the probability that all possible single-step mutations that can increase both resistance and fitness occurred in each population is very likely. The few mutations selected here as well as their repeated order with increasing CIP concentrations demonstrates that these are the most fit mutations in this *A. baumannii* strain and set of environmental conditions. The prevalence of some of these mutations in clinical samples suggests that they too may have been exposed to selection in similar conditions. Likewise, the absence of other mutations reported in shotgun mutant screens of resistance in *A. baumannii* (Edward Geisinger et al., 2018) means that these mutants produced less resistance, lower fitness, or both. Evolution experiments hold promise for ultimately forecasting mutations selected by different antimicrobials and anticipating treatment outcomes, including the diversification of the pathogen population and the likelihood of collateral sensitivity or cross-resistance (Brockhurst et al., 2019). Furthermore, knowledge of the prevailing lifestyle of the pathogen population may be critically important for treatment design. Most infections are likely caused by surface-attached populations (Wolcott, 2017; Wolcott et al., 2010), and some treatments include cycling antibiotics that promote biofilm as a primary response. For example, tobramycin is used for treating *P. aeruginosa* in cystic fibrosis patients (Hamed & Debonnett, 2017) and promotes biofilm formation (Hoffman et al., 2005; Linares, Gustafsson, Baquero, & Martinez, 2006), wherein the evolution of antibiotic resistance without a detectable fitness cost may arise during treatment. But the more diverse biofilm-adapted lineages in our experiments revealed a striking vulnerability to cephalosporins, which could provide a new strategy for treatment. Broader still, conventional wisdom has long held that the relationship between resistance and fitness is antagonistic, and that the efficacy of many antimicrobials is aided by a severe fitness cost of resistance (M. Baym et al., 2016; Hughes & Andersson, 2017; Vogwill & MacLean, 2015). This study demonstrates that the form of the relationship between fitness and resistance can be altered by the mode of growth, whereby biofilms can align resistance and fitness traits. Continued efforts to determine how the fitness landscape of various resistance pathways depends on the environment and its structure, including growth mode, could produce a valuable forecasting tool to stem the rising AMR tide.

## Methods

### Experimental evolution

Before the start of the antibiotic evolution experiment, we propagated well mixed tubes founded by one clone of the susceptible *A. baumannii* strain ATCC 17978-mff (Baumann et al., 1968; Piechaud & Second, 1951) in a modified M9 medium (referred to as M9^+^) containing 0.1 mM CaCl_2_, 1 mM MgSO_4_, 42.2 mM Na_2_HPO_4_, 22 mM KH2PO_4_, 21.7 mM NaCl, 18.7 mM NH_4_Cl and 11.1 mM glucose and supplemented with 20 mL/L MEM essential amino acids (Gibco 11130051), 10 mL/L MEM nonessential amino acids (Gibco 11140050), and 1 mL each of trace mineral solutions A, B, and C (Corning 25021-3Cl). This preadaptation phase was conducted in the absence of antibiotics for 10 days (*ca.* 66 generations) with a dilution factor of 100 per day.

After the ten days of preadaptation to M9^+^ medium, we selected a single clone and propagated for 24 hours in M9^+^ in the absence of antibiotic. We then subcultured this population into twenty replicate populations. Ten of the populations (5 planktonic and 5 biofilm) were propagated every 24 hours in constant subinhibitory concentrations of CIP, 0.0625 mg/L, which corresponds to 0.5x the minimum inhibitory concentration (MIC). After 72 hours under subinhibitory concentrations of CIP, the populations were exposed to two different antibiotic regimes for 9 more days, either constant subinhibitory concentrations of CIP or increasing concentrations of CIP (called the evolutionary rescue). For the latter, we doubled the CIP concentrations every 72 hours until 4x MIC. As a control, the 20 remaining populations were propagated in the absence of CIP (Figure 1).

We propagated the populations into fresh media every 24 hours as described by Turner *et al.* 2018 (Turner et al., 2018). For planktonic populations, we transferred a 1:100 (50 µl into 5 mL of M9^+^) dilution, which corresponded to 6.64 generations per day. For biofilm populations, we transferred a polystyrene bead (Polysciences, Inc., Warrington, PA) to fresh media containing three sterile beads. We rinsed each bead in PBS before the transfer, therefore reducing the transfer of planktonic cells. Each day we alternated between black and white marked beads, ensuring that the bacteria were growing on the bead for 24 hours, which corresponds to approximately 6 to 7.5 generations/day (Traverse et al., 2013; Turner et al., 2018). For the experiment with increasing concentrations of antibiotics, we froze a sample of each bacterial population on days 1, 3, 4, 6, 7, 9, 10 and 12. In the experiment with constant exposure to subinhibitory concentrations of antibiotics, we froze the populations on days 1, 3, 4, 9, and 12. We froze the control populations at days 1, 4, 9, and 12. For planktonic populations, we froze 1 mL of culture with 9% of DMSO. For freezing the biofilm populations, we sonicated the beads in 1 mL of PBS with a probe sonicator and subsequently froze with 9% DMSO.

### Phenotypic characterization: antimicrobial susceptibility and biofilm formation

We determined the MIC of CIP by broth microdilution in M9^+^ according to the Clinical and Laboratory Standards Institute guidelines (CLSI, 2007), in which each bacterial sample was tested to 2-fold-increasing concentration of CIP from 0.0625 to 64 mg/L. To obtain a general picture of the resistance profiles we determined the MIC to 23 antibiotics (amikacin, ampicillin, ampicillin/sulbactam, aztreonam, cefazolin, cefepime, cephalothin, meropenem, ertapenem, cefuroxime, gentamicin, CIP, piperacillin/tazobactam, cefoxitin, trimethoprim/sulfamethoxazole, cefpodoxime, ceftazidime, tobramycin, tigecycline, ticarcillin/clavulanic acid, ceftriaxone and tetracycline) by broth microdilution in commercial microtiter plates following the instructions provided by the manufacturers (Sensititre GN3F, Trek Diagnostics Inc., Westlake, OH). We tested the MIC at days 1, 3, 4, 6, 7, 9, 10 and 12 for the populations propagated under increasing concentrations of antibiotic, and at days 1 and 12 for the subinhibitory and non-antibiotic treatments. For the CIP-MICs, we used *Pseudomonas aeruginosa* PAO1 in Mueller Hinton broth as a control. No differences in the MICs were found between Mueller Hinton and M9^+^ or if measuring the MIC in 96 well-plate or in 5 mL tubes, which are the experimental conditions. Each MIC was performed in triplicate. The CIP was provided by Alfa Aesar (Alfa Aesar, Wardhill, MA). We also determined the MIC of CIP in biofilm conditions adapting the method described by Diez-Aguilar to the bead model (Díez-Aguilar et al., 2018). We resuspended each clone into fresh M9^+^ containing sterile beads (as in the experimental evolution conditions, each tube used contained three sterile beads and 5 mL of M9^+^). After 24 hours growing at 37°, each bead was propagated into new fresh M9^+^ containing different CIP concentrations (from 4 to 128 mg/L in 2-fold-increasing manner). After 24 growing at 37°, we rinsed each bead in PBS and sonicate them individually as explained before. 10 µl of the sonicated liquid were transferred to 100 µL of M9^+^. The MIC was calculated after measuring the optical density at 650 nm before and after 24 hours incubation. The inhibition of growth was defined as the lowest antibiotic that resulted in an OD difference at or below 0.05 after 6 hours of incubation.

We estimated the biofilm formation of the selected clones using a modification of the previously described protocol (O’Toole & Kolter, 1998). We resurrected each clone in 5 mL of M9^+^ containing 0.5 mg/L of CIP and grew them for 24 hours. For each strain, we transferred 50 µl into 15 mL of M9^+^. We tested 200 µl of the previous dilution of each clone to 4 different subinhibitory CIP concentrations (0 mg/L, 0.01 mg/L, 0.03 mg/L and 0.0625 mg/L). After 24 hours of growing at 37°C, we measured population sizes by optical density (OD) at 590nm (OD_Populations_). Then, we added 250 µl of 0.1% crystal violet and incubated at room temperature for 15 minutes. After washing the wells and drying for 24 hours, we added 250 µl 95% EtOH solution (95% EtOH, 4.95% dH2O, 0.05% Triton X-100) to each well and incubated for 15 minutes and biofilm formation was measured by the OD at 590nm (OD_Biofilm_). Biofilm formation was corrected by population sizes (OD_Biofilm_/OD_Population_). Results are the average of three experiments (Figure S3).

### Fitness measurement

We selected 5 biofilm and 5 planktonic clones at the end of the evolutionary rescue experiment (Figure 2) and determined the fitness by directly competing the ancestral strain and the evolved clone variants both in planktonic and in biofilm conditions (Figure 3) (Turner et al., 2018). We revived each clone from a freezer stock in M9^+^ for 24 hours. We maintained the same evolutionary conditions to revive the clones, adding 3 beads and/or CIP to the broth when required. After 24 hours, we added equal volume of the clones and the ancestors in M9^+^ in the absence of antibiotics. For planktonic populations, we mixed 25 µl of each competitor in 5 mL of M9^+^. For biofilm competitions, we sonicated one bead per competitor in 1 mL of PBS and mixed in 5 mL of M9^+^ containing 3 beads. The mix was cultured at 37°C for 24 hours. We plated at time zero and after 24 hours. For each competition, we plated aliquots onto nonselective tryptic soy agar and tryptic soy agar containing CIP. Selection rate (*r*) was calculated as the difference of the Malthusian parameters for the two competitors: *r* = (ln(CIP resistant_d=1_/CIP resistant_d=0_))/(ln(CIP susceptible_d=1_/CIP susceptible_d=0_))/day (Lenski, 1991). Susceptible populations were calculated as the difference between the total populations (number of colonies/mL growing on the nonselective plates) and the resistant fraction (number of colonies/mL growing on the plates containing CIP). As a control for calculating the correct ratio of susceptible vs. resistant populations, we replica plated 50 to 100 colonies from the nonselective plates onto plates containing CIP as previously described (Santos-Lopez et al., 2017). Results are the average of three to five independent experiments.

### Genome sequencing

We sequenced whole populations of three evolving replicates per treatment. We sequenced the populations at days 1, 3, 4, 6, 7, 9, 10, and 12 of the populations under increasing concentrations of CIP (populations P1, P2, P3 and B1, B2, B3 for planktonic and biofilm populations) and at days 1, 4, 9, and 12 of the populations under subinhibitory concentration and no antibiotic treatments. In addition, we selected 49 clones for sequencing at the end of the experiment (Figure 2F). 12 of the clones were recovered from the populations propagated in the absence of the antibiotic, 12 clones from the subinhibitory concentrations of CIP treatment and 25 were isolated from the increasing concentrations of antibiotic. We revived each population or clone from a freezer stock in the growth conditions under which they were isolated (*i.e.* the same CIP concentration which they were exposed to during the experiment) and grew for 24 hours. DNA was extracted using the Qiagen DNAeasy Blood and Tissue kit (Qiagen, Hiden, Germany). The sequencing library was prepared as described by Turner and colleagues (Turner et al., 2018) according to the protocol of Baym *et al.* (Baym et al., 2015), using the Illumina Nextera kit (Illumina Inc., San Diego, CA) and sequenced using an Illumina NextSeq500 at the Microbial Genome Sequencing center (http://micropopbio.org/sequencing.html).

### Data processing

All sequences were first quality filtered and trimmed with the Trimmomatic software v0.36 (Bolger, Lohse, & Usadel, 2014) using the criteria: LEADING:20 TRAILING:20 SLIDINGWINDOW:4:20 MINLEN:70. Variants were called with the breseq software v0.31.0 (Deatherage & Barrick, 2014) using the default parameters and the -p flag when required for identifying polymorphisms in populations. This option calls a mutation if it is observed in 2 reads from each strand and reaches 5% in the population. The average depth of sequencing for populations was 219 + 51x and average genome coverage was 98.7 + 0.128%. The reference genome used for variant calling was downloaded from the NCBI RefSeq database using the 17-Mar-2017 version of *A. baumannii* ATCC 17978-mff complete genome (GCF_001077675.1). In addition to the chromosome NZ_CP012004 and plasmid NZ_CP012005 sequences, we added two additional plasmid sequences to the reference genome that are known to be present in our working strain of *A. baumannii* ATCC 17978-mff: NC009083, NC_009084. Mutations were then manually curated and filtered to remove false positives. Mutations were filtered if the gene was found to contain a mutation when the ancestor sequence was compared to the reference genome or if a mutation never reached a cumulative frequency of 10% across all replicate populations. Diversity measurements were made in R using the Shannon index considering the presence, absence, and frequency of alleles. Significant differences between biofilm and planktonic populations were determined by the Kruskal Wallis test. Filtering, mutational dynamics, and plotting were done in R v3.4.4 (www.r-project.org) with the packages ggplot2 v2.2.1 (Wickham, 2016), dplyr v0.7.4 (Wickham, François, Henry, & Müller, 2018), vegan v2.5-1 (Oksanen et al., 2018), and reshape2 (Wickham, 2007).

### Data Availability

R code for filtering and data processing can be found here: https://github.com/sirmicrobe/U01_allele_freq_code. All sequences were deposited into NCBI under the Biosample accession numbers SAMN09783599-SAMN09783682.

## Authors contribution

VSC, AS-L and CWM conceived and designed the study; AS-L and MRS performed the experiments; DS sequenced the samples; CWM did the bioinformatic analysis; AS-L, CWM and VSC wrote the paper.

## Acknowledgments

We thank Caroline B. Turner for helpful discussions and proofreading of the paper, Allison L. Welp for laboratory assistance and Christopher Deitrick for depositing the sequences in the NCBI database. This research was supported by NIH U01AI124302-01.

---

**Figure S1.**
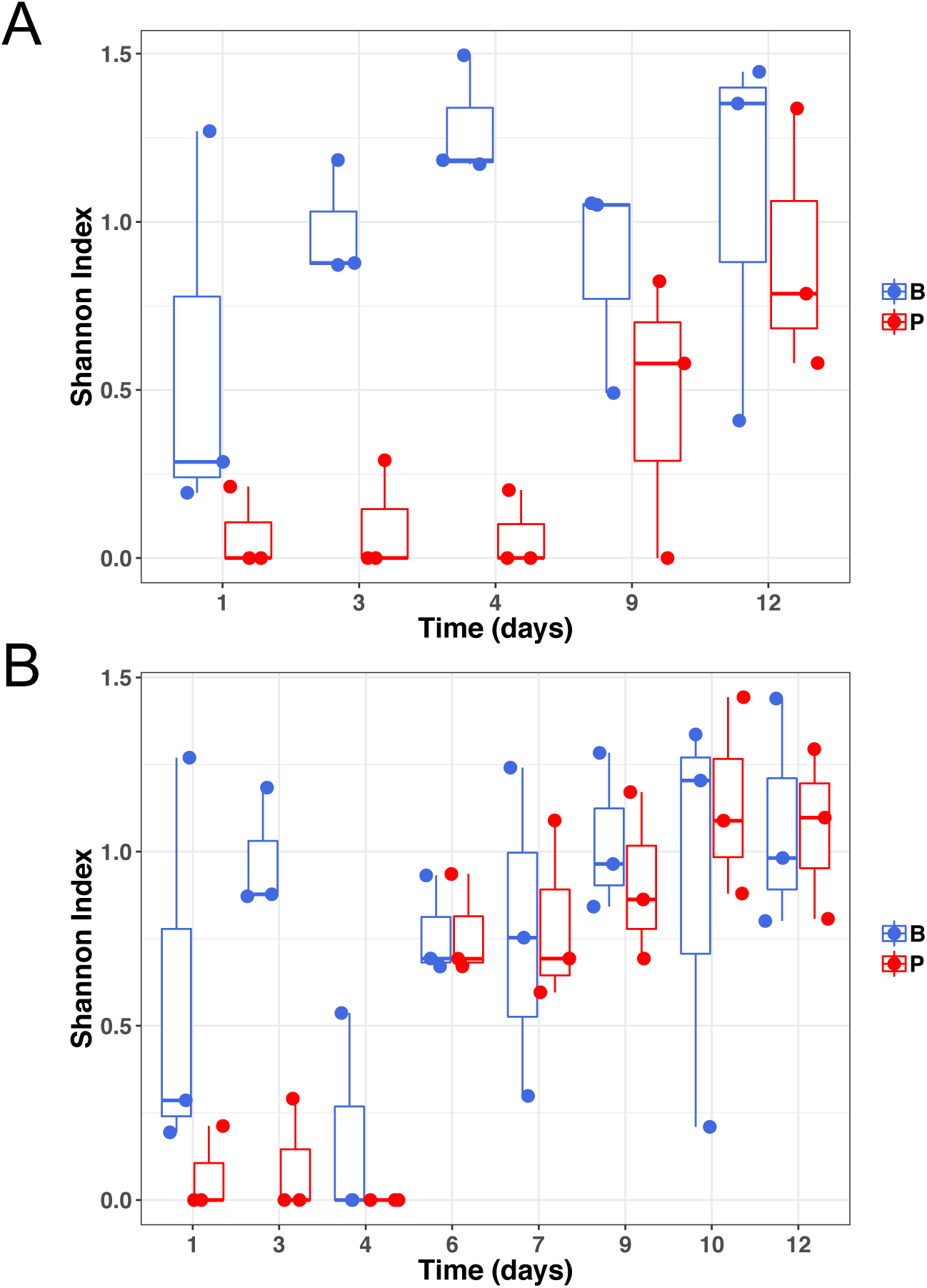
Genetic diversity of samples at subinhibitory concentrations of ciprofloxacin (A) or during the evolutionary rescue experiment with increasing concentrations of ciprofloxacin (B). Biofilm populations in blue and planktonic populations in red.

**Figure S2.**
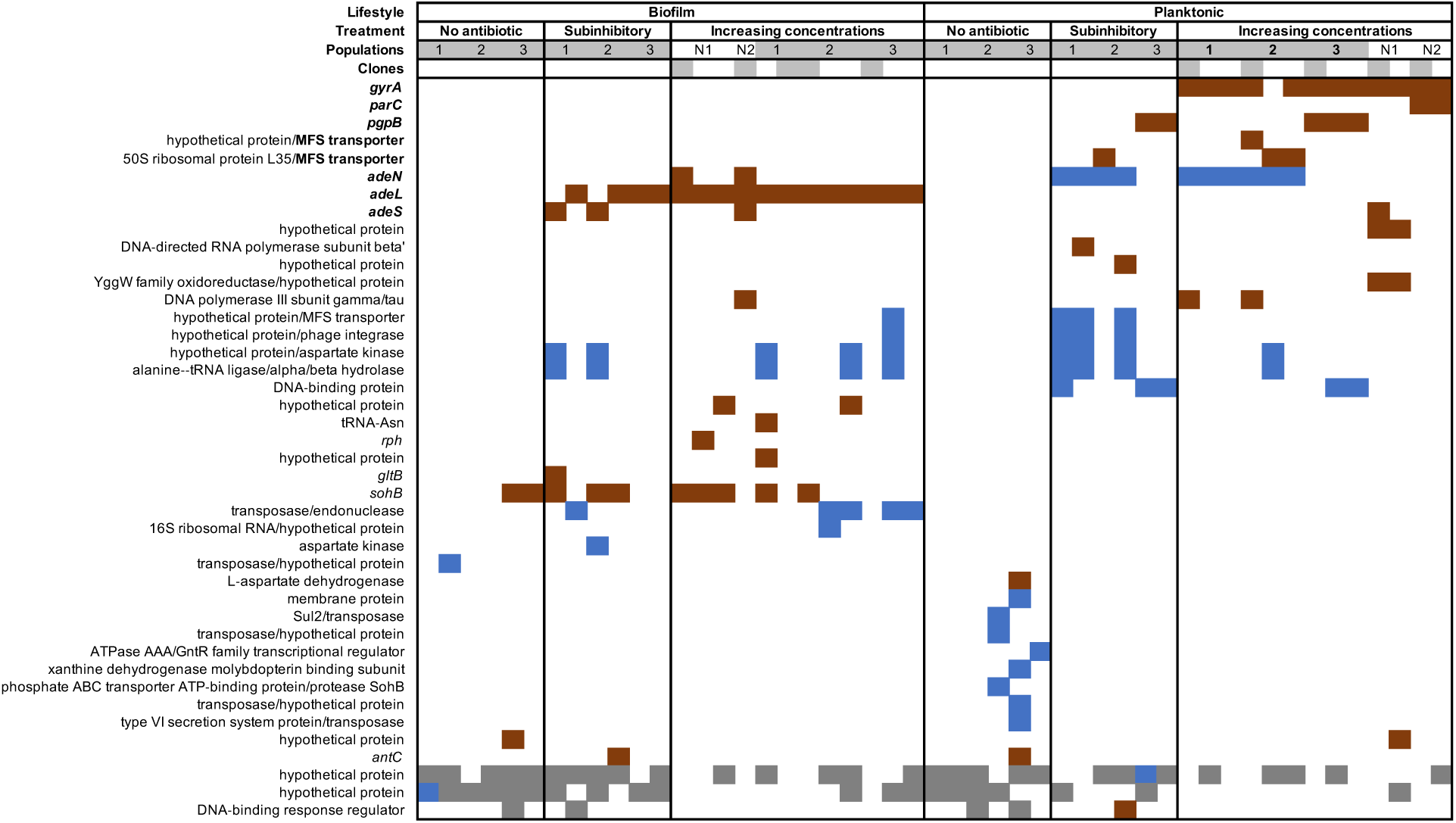
Mutated genes in the sequenced clones differ between treatments. Each column represents one clone. Grey shaded clones were used for MIC and fitness estimations. Red color indicates SNPs or small indels, blue color indicates new junction evidences and grey indicates missing coverage indicative of a deletion.

**Figure S3.**
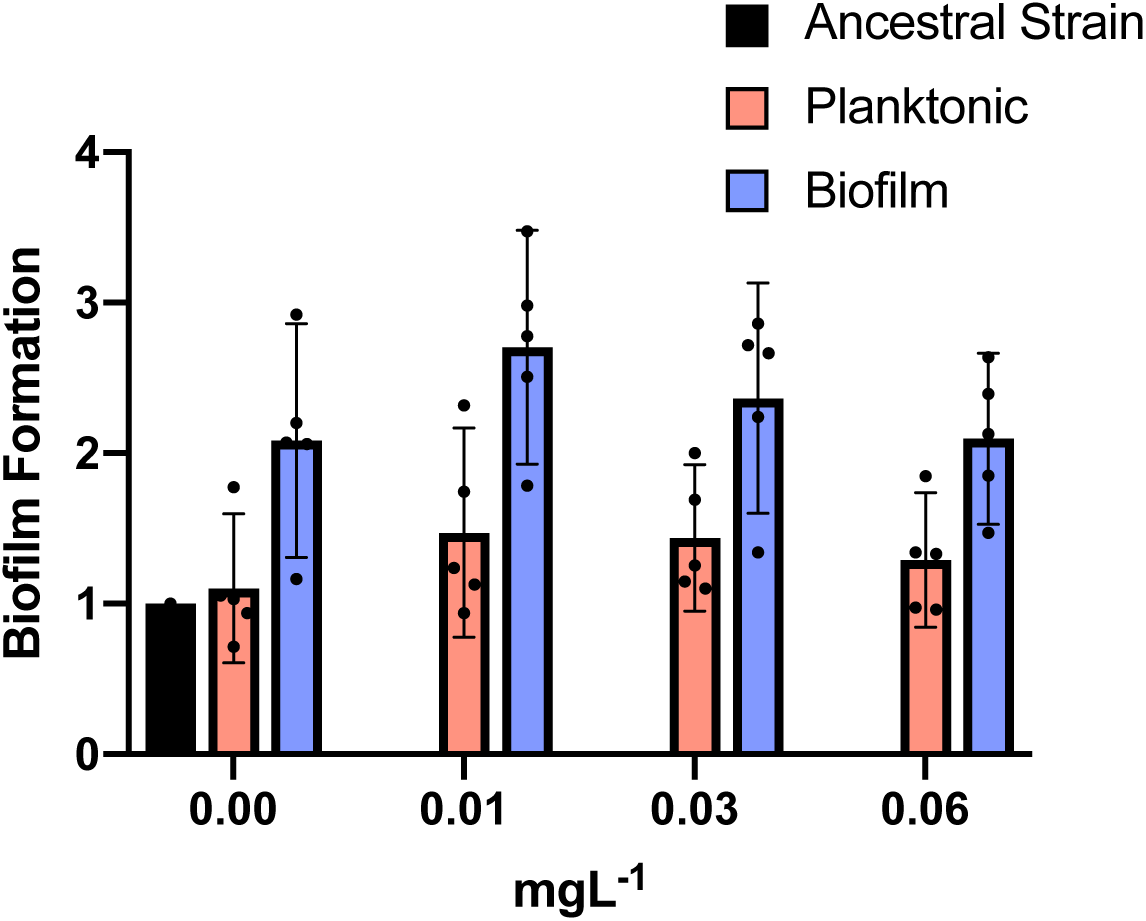
Biofilm production under subinhibitory concentrations of CIP. Blue and red bars show biofilm and planktonic clones, respectively. The ancestral strain is represented by the black bar. Individual clone results are shown as points. The averages are shown by bars. 95% CI are indicated by the error bars. Biofilm clones produced more biofilm than planktonic clones: two tailed t-test of biofilm formation with 0.00 mg/L of CIP: p = 0.006, t = 3.008, df =32; with 0.01 mg/L of CIP: p = 0.0006, t = 3.780, df =32; with 0.03 mg/L of CIP: p = 0.0077 t = 2.841, df = 32 and with 0.0625 mg/L of CIP: p = 0.018 t = 2.471 df = 32.

**Table S1.**
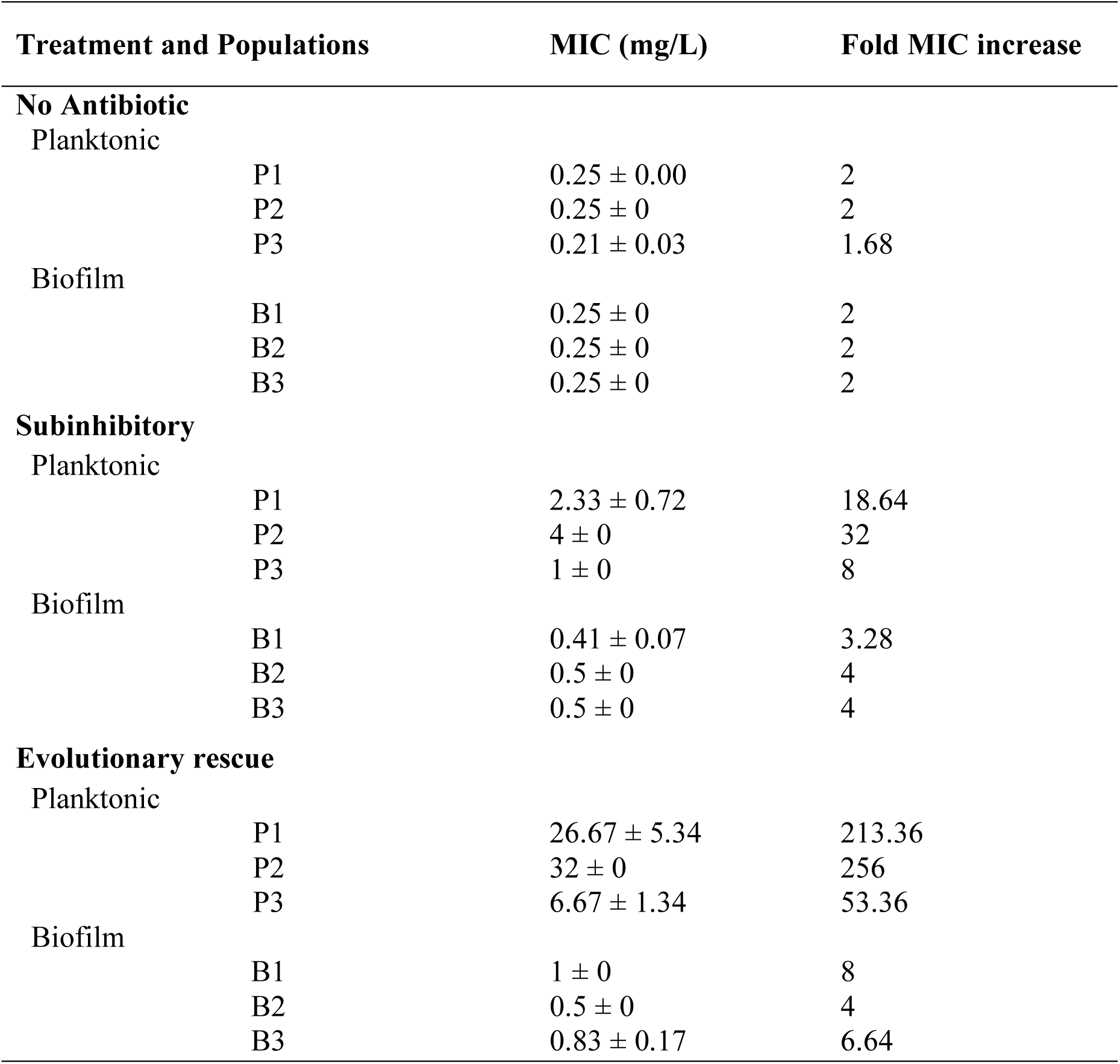
Antibiotic susceptibility of the populations propagated in the absence, in subinhibitory concentrations or increasing concentrations of CIP at the end of the experiment (day 12). MICs are expressed in mg/L and standard errors of the mean are indicated. Fold increase in MIC compared to the ancestral strain are also indicated.

| Treatment and Populations |  | MIC (mg/L) | Fold MIC increase |
| --- | --- | --- | --- |
| **No Antibiotic** |  |  |  |
| Planktonic | P1 | $0.25 \pm 0.00$ | 2 |
| | P2 | $0.25 \pm 0$ | 2 |
| | P3 | $0.21 \pm 0.03$ | 1.68 |
| Biofilm | B1 | $0.25 \pm 0$ | 2 |
| | B2 | $0.25 \pm 0$ | 2 |
| | B3 | $0.25 \pm 0$ | 2 |
| **Subinhibitory** |  |  |  |
| Planktonic | P1 | $2.33 \pm 0.72$ | 18.64 |
| | P2 | $4 \pm 0$ | 32 |
| | P3 | $1 \pm 0$ | 8 |
| Biofilm | B1 | $0.41 \pm 0.07$ | 3.28 |
| | B2 | $0.5 \pm 0$ | 4 |
| | B3 | $0.5 \pm 0$ | 4 |
| **Evolutionary rescue** |  |  |  |
| Planktonic | P1 | $26.67 \pm 5.34$ | 213.36 |
| | P2 | $32 \pm 0$ | 256 |
| | P3 | $6.67 \pm 1.34$ | 53.36 |
| Biofilm | B1 | $1 \pm 0$ | 8 |
| | B2 | $0.5 \pm 0$ | 4 |
| | B3 | $0.83 \pm 0.17$ | 6.64 |

**Table S2.**
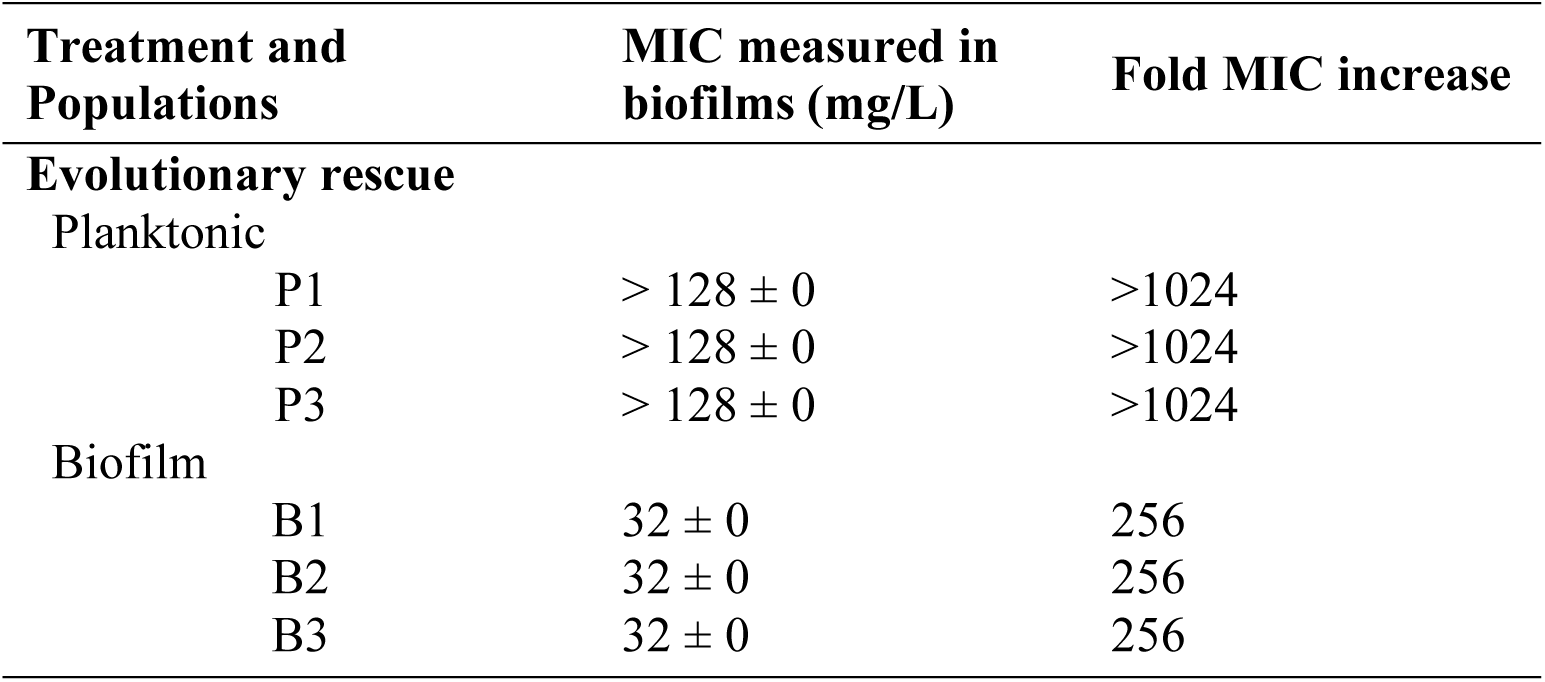
Antibiotic susceptibility of one clone of each population propagated in increasing concentrations of CIP at the end of the experiment (day 12). MICs were measured in biofilms and are expressed in mg/L and standard errors of the mean are indicated. Fold increase in MIC compared to the ancestral strain are also indicated.

**Table S3. Mutation probabilities (attached XLS file)**

**Table S4.**
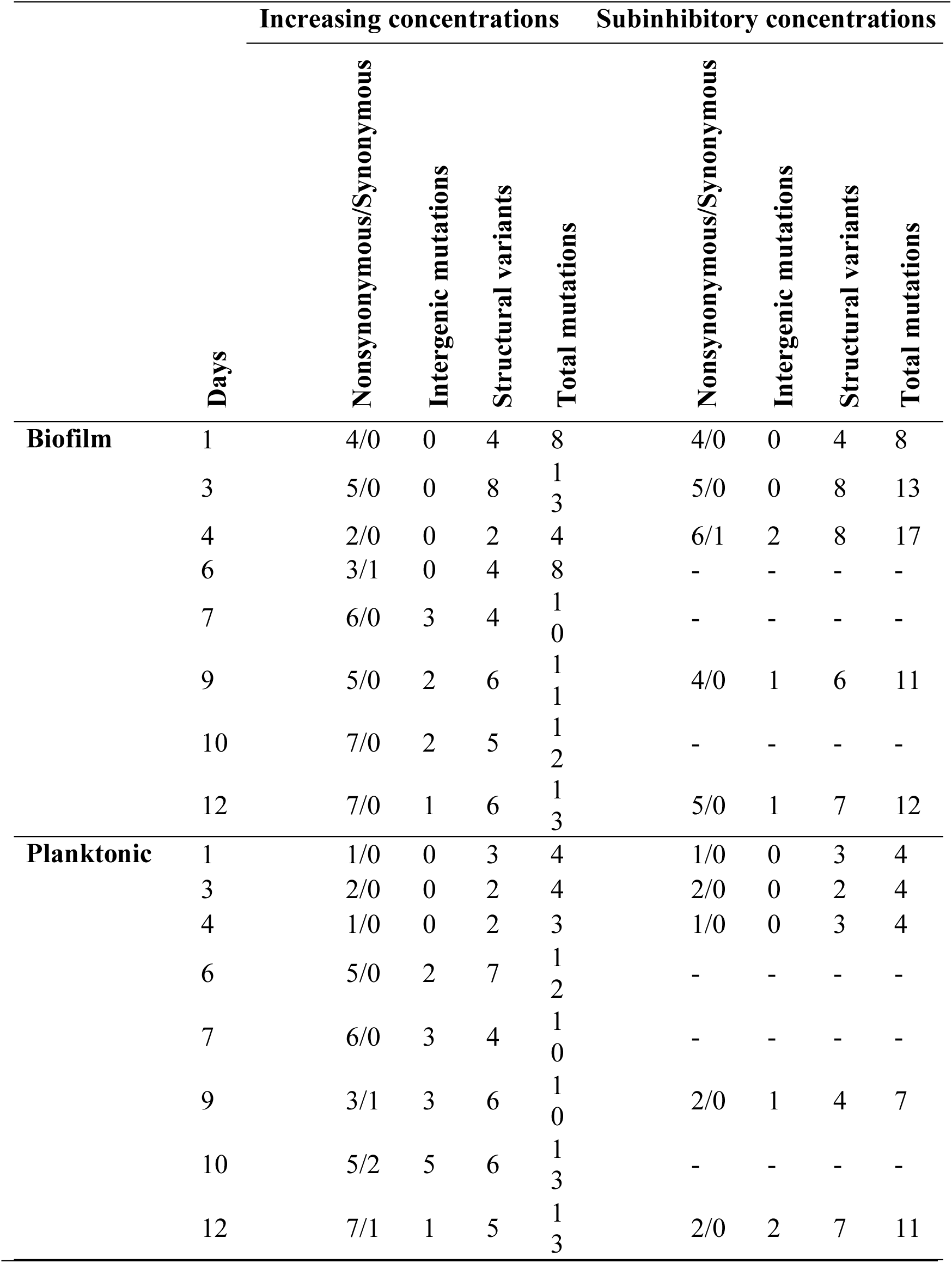
Attributes of the contending mutations each day in the experimental evolution.

**Table S5.**
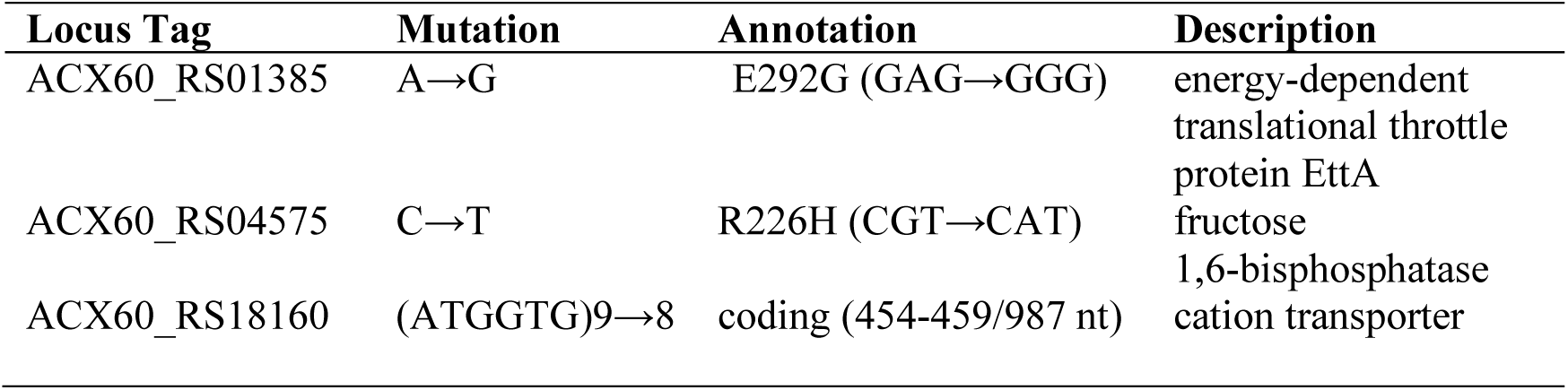
Mutated genes in the ancestral strain compared to the *A. baumannii* ATCC 17978-mff complete genome (GCF_001077675.1) after 10 days of evolution in M9^+^.

| **Locus Tag** | **Mutation** | **Annotation** | **Description** |
| --- | --- | --- | --- |
| ACX60_RS01385 | A→G | E292G (GAG→GGG) | energy-dependent translational throttle protein EttA |
| ACX60_RS04575 | C→T | R226H (CGT→CAT) | fructose 1,6-bisphosphatase |
| ACX60_RS18160 | (ATGGTG)9→8 | coding (454-459/987 nt) | cation transporter |

**Table S6. Complete list of mutated genes from the sequenced clones (attached XLS file)**

**Table S7.**
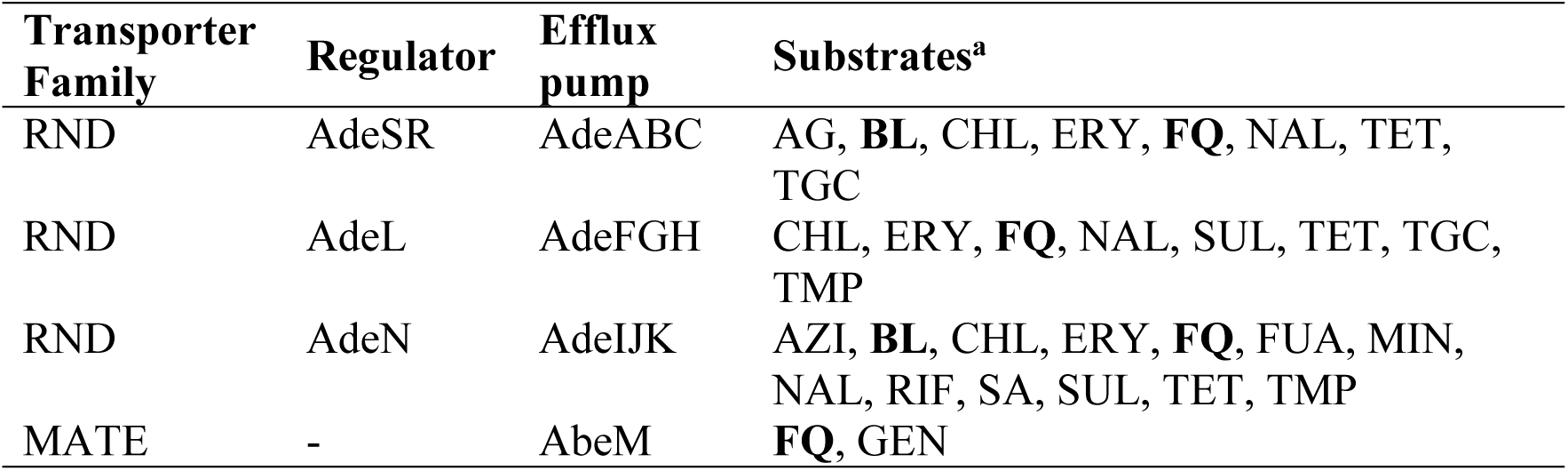
Efflux pumps and their regulators in *A. baumannii* 17978 targeted under CIP pressure. Table adapted from (Li, Elkins et al. 2016). AG aminoglycosides, AZI azithromycin, BL β-lactams, CHL chloramphenicol, CIP CIP, CL clindamycin, ERY erythromycin, FLO florfenicol, FUA fusidic acid, GEN gentamicin, MIN minocycline, NAL nalidixic acid, NOR norfloxacin, RIF rifampicin, SUL sulfonamides, TET tetracycline, TGC tigecycline, TMP trimethoprim. ^a^ References in (X.-Z. Li et al., 2016).

